# Protein structure prediction using sparse NOE and RDC restraints with Rosetta in CASP13

**DOI:** 10.1101/597724

**Authors:** Georg Kuenze, Jens Meiler

## Abstract

Computational methods that produce accurate protein structure models from limited experimental data, e.g. from nuclear magnetic resonance (NMR) spectroscopy, hold great potential for biomedical research. The NMR-assisted modeling challenge in CASP13 provided a blind test to explore the capabilities and limitations of current modeling techniques in leveraging NMR data which had high sparsity, ambiguity and error rate for protein structure prediction. We describe our approach to predict the structure of these proteins leveraging the Rosetta software suite. Protein structure models were predicted *de novo* using a two-stage protocol. First, low-resolution models were generated with the Rosetta *de novo* method guided by non-ambiguous nuclear Overhauser effect (NOE) contacts and residual dipolar coupling (RDC) restraints. Second, iterative model hybridization and fragment insertion with the Rosetta comparative modeling method was used to refine and regularize models guided by all ambiguous and non-ambiguous NOE contacts and RDCs. Nine out of 16 of the Rosetta *de novo* models had the correct fold (GDT-TS score >45) and in three cases high-resolution models were achieved (RMSD <3.5 Å). We also show that a meta-approach applying iterative Rosetta+NMR refinement on server-predicted models which employed non-NMR-contacts and structural templates leads to substantial improvement in model quality. Integrating these data-assisted refinement strategies with innovative non-data-assisted approaches which became possible in CASP13 such as high precision contact prediction will in the near future enable structure determination for large proteins that are outside of the realm of conventional NMR.

## Introduction

A current focus in structural biology has been the development of advanced integrative modeling techniques that can determine structures of proteins and their interactions from limited experimental data^1,2^. Those methods are called in when classical structural biology techniques such as X-ray crystallography and nuclear magnetic resonance (NMR) spectroscopy fail to obtain complete and unambiguous data at atomic-detail. NMR can obtain such data for small proteins under physiologically conditions but loses quickly resolution and sensitivity when the protein under study becomes large (>20 kDa). The classical approach of collecting short-range inter-proton distance measurements by nuclear Overhauser effect spectroscopy (NOESY) becomes difficult due to peak line broadening and low signal-to-noise ratio. Low-resolution and sparse NMR datasets thus call for computational methods that can translate them into accurate structural models. At the same time these methods add atomic-detail information to the model which may not be present in the NMR data, e.g. sidechain positions.

The Rosetta program^3^ offers a unique platform of integrative modeling tools and has been designed to make use of different types of NMR data. For example, chemical shifts (CSs)^4-6^ and residual dipolar couplings (RDCs)^7^ can be used to guide the search and assembly of small peptide fragments with known conformations from which Rosetta builds a protein structure *de novo*. Using only this kind of backbone NMR data, which is available at an early stage of the NMR structure determination process, this method, called ‘CS-Rosetta’, was able to correctly model the structure of proteins up to 25 kDa^8^. Incorporation of sparse NOEs from selectively ILV-labeled deuterated proteins and improvements to the conformational sampling algorithm were shown to increase the application limit to 40 kDa^9^. Moreover, the CS-Rosetta method was extended to include sparse contact information^10^ and structural templates^11^ from homologous proteins with guidance from chemical shift-based alignments. With the impressive advancement in coevolution- and deep-learning-based contact prediction methods^12^, evolutionary couplings (ECs) become now increasingly available as new type of distance restraints for Rosetta modeling to supplement sparse NOEs^13,14^.

The aforementioned CS-Rosetta studies used expert-collected experimental NMR datasets with high completeness in the sense that chemical shift assignments were available for almost all residues. In addition, two or more RDC datasets and one to two NOEs per residue were used^9^. The data provided in CASP13 comprised simulated data and one real NMR dataset. The number of NOE restraints was comparable to that one in previous CS-Rosetta studies but datasets contained a considerable number of NOEs with high ambiguity as well as incorrectly assigned NOEs. Furthermore, in order to simulate realistic difficulties in the data collection process, e.g. line broadening due to internal protein motions, residues were removed from the peak list, and hence, the assignment became incomplete; RDCs, CS-derived torsion angle restraints and NOEs were available for only part of the protein. It is an interesting question how sparse and ambiguous NMR data are best incorporated into modeling methods, to which extent they can improve the accuracy of the predicted structural model and whether they can provide higher accuracy than so-called ‘free modeling’ techniques which omit the use of experimental data.

In CASP13, we employed a two-stage approach adopted to the ambiguity level of the restraints: initial fold-level modeling guided by unambiguous data followed by iterative model hybridization and refinement using all NMR data. Our results show that with this low amount of NMR information Rosetta can generate models that have the correct fold and are in some cases very close to the native structure. In our analysis after CASP13, we further explored the possibility of combining NMR-assisted modeling with state-of-the-art free modeling techniques which have seen considerable improvements in the last CASP assessment. The main driving forces seem to be the increasing availability of structural templates^15-17^ and the high precision of contact predictions which have become possible due to new technologies like deep convolutional neural networks that allow efficient use of coevolution information^18-21^. As those new modeling techniques are made available to users, e.g. in the form of a public webserver, they can be easily incorporated into the NMR structure prediction protocol. Here, we demonstrate that models submitted by modeling servers can be easily recombined and hybridized with Rosetta and refined with NMR data to yield structure predictions with better accuracy than Rosetta-NMR models and the original server predictions. We therefore suggest a ‘meta-approach’ to NMR structure prediction which supplements sparse experimental data with complementary restraints, e.g. from homolog templates and predicted contacts. This meta-approach can be a constructive way to determine the structures of challenging protein targets which till now were outside of the realm of solution state NMR.

## Materials and Methods

### Overall structure prediction protocol

As in previous CASP contact-assisted experiments^22,23^, a two-stage modeling approach consisting of initial fold-level modeling and subsequent model refinement was employed (**Figure 1**).

**Figure 1.**
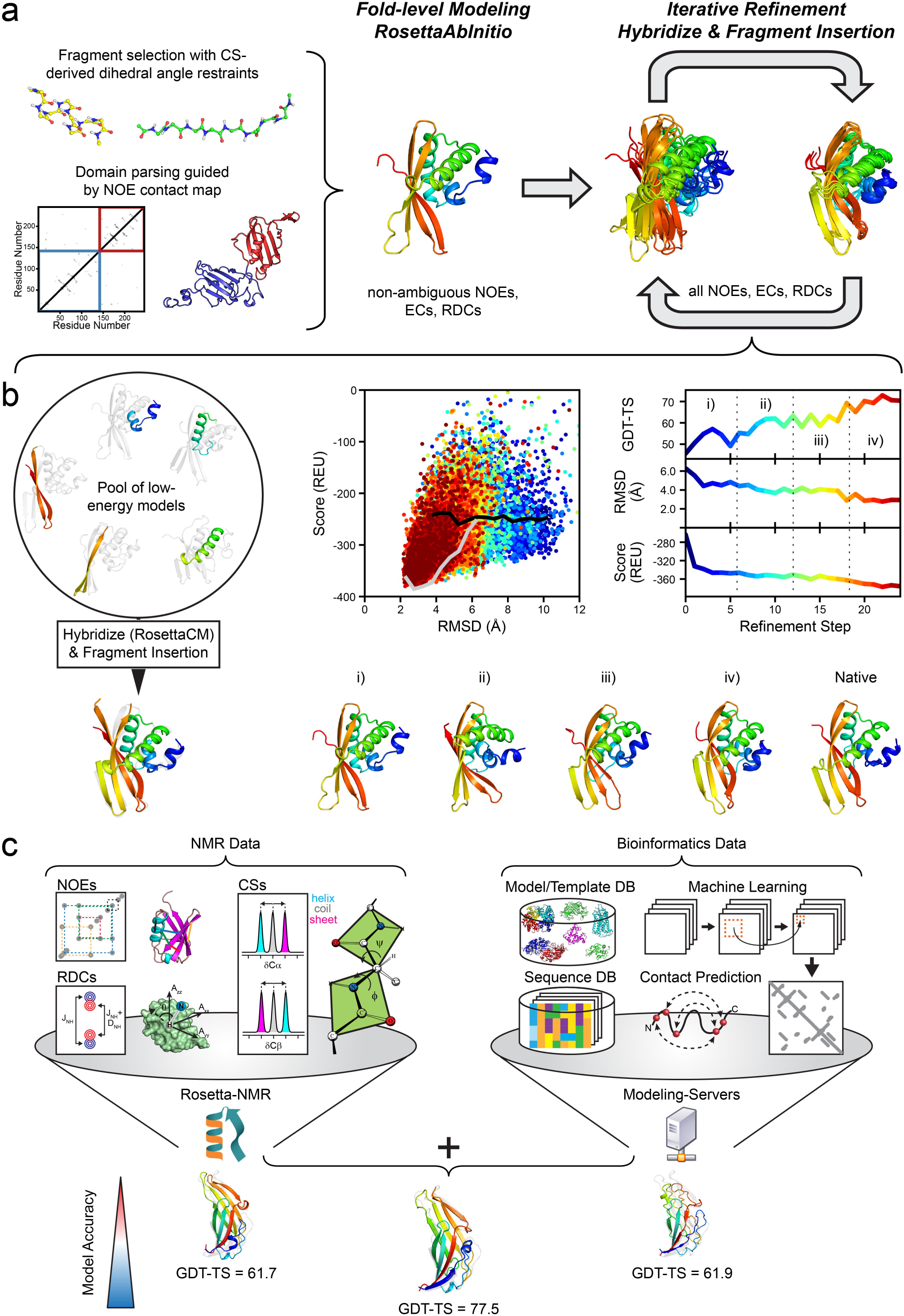
Overview of the modeling protocol employed in the CASP13 NMR-assisted structure prediction category. **(a)** Overall flowchart: Initial fold-level modeling was done with RosettaAbInitio guided by non-ambiguous NOEs, evolutionary coupling restraints (ECs) and RDCs. The library of 3mer and 9mer peptide fragments was created with the Rosetta Fragment Picker using CS-derived □/ψ dihedral angle restraints. Proteins with >200 residues were manually parsed into domains guided by non-ambiguous NOE contacts, and domains were modeled separately. After fold-level modeling, models were iteratively refined by hybridization and fragment insertion guided by all ambiguous and non-ambiguous NOEs, ECs and RDCs. **(b)** Improvement in model accuracy for target N0968s1 through an iterative refinement protocol. The protocol maintains a pool of low-energy models which are hybridized with RosettaCM and diversified through fragment insertion. The protocol stops when models are converged in terms of their pairwise GDT-HA or the final refinement step is reached. The bottom row demonstrates the improvement in the accuracy of N0968s1 models as the protocol proceeds: (i) GDT-TS = 47.5, (ii) GDT-TS = 58.0, (iii) GDT-TS = 60.2, (iv) GDT-TS = 72.4. **(c)** Suggested meta-approach to NMR structure modeling by incorporating structural restraints retrieved from bioinformatical sources/databases (DB), e.g. predicted residue-residue contacts and structural templates. In our post-CASP13 analysis, we used the initial predictions of five different servers (Robetta^34^, I-TASSER^35,36^, QUARK^37^, RaptorX-Contact^38^, RaptorX-TBM^39^) and refined them iteratively with Rosetta and NMR data leading to more accurate models than each individual technique. Model improvement is exemplified for target N0981-D5 for which the GDT-TS score increased by >15.

The provided NMR data comprised simulated ^1^H-^1^H-NOE and ^1^H-^15^N-RDC data for 11 out of 12 protein targets as well as one real experimental ^1^H-^1^H-NOE dataset for one protein target. In addition, □/ψ dihedral angle ranges (with ±30° uncertainty) which had been back-calculated from the simulated or real chemical shifts (CSs) were provided by the organizers. The NMR data were sparse and ambiguous, i.e. many NOE cross-peaks could be assigned to more than one possible pair of H-atoms. In addition, the NOE dataset contained a considerable fraction of false positive (FP) contacts arising from incorrect peak assignments because NOEs had been extracted from NOESY-NMR spectra simulated with realistic peak line widths and signal-to-noise ratio. Thus, denoising and optimal use of the ambiguous data along the modeling pipeline was important.

In the first modeling stage, only non-ambiguous NOEs and RDCs were used. In the following refinement stage, starting from these fold-level models, ambiguous data were included (**Figure 1a**). These NMR data were further supplemented by evolutionary coupling (EC) distance restraints which were provided by the CASP organizers and had been computed with the MetaPSICOV method^24^.

Parallel to calculations with this full set of restraints (i.e. dihedral, NOE, RDC and EC restraints), a second independent prediction using only dihedral and RDC restraints was made for every target, except N1008 for which no RDCs had been measured. Purpose of this minimal restraint set was to test the efficacy of backbone-only data for NMR structure prediction. The best-scoring model made with the full set of restraints was submitted as Model 1, whereas the RDC-only model was submitted as one of the remaining four models, usually Model 2.

Proteins larger than 200 residues in length for which domain boundaries could be unambiguously identified based on the NOE contact map were parsed manually into domains. These domains were modeled and refined separately, and afterwards recombined to the full-length model with RosettaCM^25^. This strategy was applied to targets N0989 (246 residues) and N1005 (326 residues). Similarly, the heterodimeric target N0980 was predicted by first modeling and refining its two subunits (N0980s1 and N0980s2) separately, and afterwards assembling them with RosettaDock^26^.

### Fold-level modeling

The Rosetta *de novo* structure prediction protocol^27,28^, referred to as RosettaAbInitio, was used for initial fold-level modeling (**Figure 1a**) generating 10,000 models for each target. The fragment library for the RosettaAbInitio protocol contained 200 3mer and 9mer peptide fragments per residue position which were selected with the Rosetta Fragment Picker^29^ guided by PSIPRED^30^ and Jufo9D^31^ secondary structure predictions and the CS-derived dihedral angle restraints. Non-ambiguous NOEs, ECs and RDCs were included in the scoring function with weights that were adjusted such that the sum of the restraint scores was approximately equal to the Rosetta energy.

After fragment assembly, ‘centroid’ (i.e. coarse-grained representation in Rosetta) models were converted to all-atom models and subjected to a short optimization in both internal and cartesian space using the RosettaFastRelax^32^ protocol.

One hundred models with the lowest combined Rosetta energy and restraint score were selected for the subsequent refinement stage. In order to maintain structural diversity in the model pool a minimum mutual distance between models corresponding roughly to a TM-score^33^ of 0.75 was enforced in the selection step. Furthermore, a penalty was applied to models which were dissimilar to the ‘reference’ model, i.e. the lowest energy representative from the three largest model clusters, by more than 25% GDT-HA.

### Model refinement

Initial fold-level models were recombined and refined using an iterative version of the RosettaCM^25^ protocol (**Figure 1b**) which was originally developed for comparative modeling. Structural optimization was accomplished by extracting and recombining secondary structure segments from a pool of low energy models together with fragment insertion. At each refinement step, 480 to 720 new models were generated and the best 100 models were selected based on the sum of their Rosetta energy and NMR restraint score for the next iteration. The minimal mutual model distance was gradually lowered in subsequent selection steps. Refinement was continued until the model pool was converged in terms of the pairwise GDT-HA or the maximum refinement step which was possible within the time constraint of the CASP experiment was reached.

After the final refinement step, the models with the lowest Rosetta all-atom energy and restraint score were visually inspected. If convergence was reached the top-scoring model was selected and submitted as Model 1. If these models varied significantly, they were clustered. Models usually fell within two to three clusters and the models corresponding to the cluster centroids were selected for submission.

### Incorporation of server-models into Rosetta+NMR refinement

As small adjustment to the described two-stage protocol, we explored in our analysis after CASP13 whether the use of template information can improve our structure predictions (**Figure 1c**). We chose to include the submitted models of five servers which had the best performance in the previous CASP12 experiment (Robetta^34^, I-TASSER^35,36^, QUARK^37^, RaptorX-Contact^38^, RaptorX-TBM^39^) into the RosettaCM refinement stage. Those models leverage other types of restraints than NMR data which is why we consider their incorporation a ‘meta-approach’ to NMR structure prediction (**Figure 1c**). Models were recombined and refined with NMR data through 20 rounds of RosettaCM. The model with the lowest combined Rosetta energy and restraint score after the last refinement step was deemed the final model and compared to the experimental reference structure.

### Incorporation of NMR restraints

Contacts inferred from non-ambiguously assigned NOE cross-peaks were used as ‘strong’ restraints with a flat-bottom bounded penalty function and applied during all stages of the modeling protocol. Only contacts between residues which were more than five sequence positions apart were kept in order to avoid over-constraining and distorting the local model geometry.

The applied NOE penalty function grows quadratically outside of the lower (l_b_) and upper (u_b_) bound, and linearly at distances larger than 0.5 Å beyond the upper bound which was set to the simulated NOE distance plus an additional 1.5 Å padding. The lower bound was set to 1.5 Å.

In the low-resolution phase of the RosettaAbInitio and RosettaCM protocols in which the protein adopts a coarse-grained representation and the sidechain is treated by a single ‘centroid’ atom, sidechain-sidechain NOEs were mapped onto the centroid atom as described previously^9^. The upper bound of the mapped restraint was increased to u_b,map_ = u_b_ + h where h is the number of methyl groups involved in the restraint (0, 1 or 2). During full-atom modeling, sidechain-sidechain NOE restraints involving groups of equivalent or non-stereochemically assigned protons were evaluated after applying a r^-6^ distance averaging.

Ambiguous NOE contacts were incorporated in the second refinement stage: as sigmoidal restraints between Cβ atoms (Cα in case of glycine) in centroid phases and as groups of nested bounded restraints when the protein was in full-atom representation. A sigmoidal restraint was created for every residue pair with sequence separation of six or higher belonging to one or more ambiguous contacts in the NOE peak list. A weighting factor was applied for each restrained residue pair which was set inversely proportional to the ambiguity level (i.e. inverse of the group size) of the ambiguous NOE. The final weight was the sum of this ratio over all ambiguous NOEs which a particular residue pair had been assigned to. Only the highest scoring 3L/2 (L is the sequence length) restraints were used. The sigmoidal scoring function was centered at a Cβ-Cβ distance of 8 Å and offset by a value of - 0.5 such that the restraint score fell with the range from −1 (satisfied) to 0 (non-satisfied).

For full-atom modeling, ambiguous NOE contacts were represented as a group of nested bounded restraints with a penalty function set up as described above. Only the lowest scoring restraint from this group was considered in calculating the total NOE restraint score.

Like ambiguous NOEs, the MetaPSICOV-predicted EC contacts were incorporated as sigmoidal restraints between Cβ atoms (Cα in case of glycine) centered at a 8 Å distance and weighted by their confidence score. Only the L most confident restraints (L is the sequence length) with a sequence separation more than five residues were used.

Simulated amide-backbone RDCs were added as additional pseudo-energy to the restraint score. The RDC score was thereby calculated as sum of squared errors between simulated and model-predicted RDC values after computation of the molecular alignment tensor by singular value decomposition (SVD).

CS-back-calculated dihedral angle ranges were used as □/ψ angle restraints in the fragment selection process and scored with a periodic bounded penalty function which had a periodicity of 2π and grew quadratically outside of the specified dihedral angle range.

## Results

### Rosetta modeling translates limited NMR data into accurate structural models

An aim of the NMR-assisted structure prediction experiment in CASP13 was to investigate whether modeling techniques can leverage limited and ambiguous NMR data for protein structure modeling and whether NMR data improve the prediction accuracy. In order to mimic realistic conditions typically found in NMR studies of larger (>20 kDa) and dynamic proteins, the NMR datasets were sparse and contained erroneous NOEs. In addition, NMR data assignments covered only part (~50%) of the protein.

With our strategy of translating the provided NMR data into structural restraints after discarding contacts with minimal sequence separation (|i-j| ≤ 5) the average number of NOE, RDC and □/ψ restraints per residue was 2.2, 0.8 and 0.5, respectively (**Figure S1a**). These are far fewer restraints than needed for experimentally driven NMR structure calculations which typically require 40-50 NOE contacts per residue. Moreover, a considerable fraction of NOE contacts were false positives (FP-NOEs) due to assignment errors or missing NOE peaks which were accounted for in the simulation by peak line broadening and low signal-to-noise ratio. By comparison with the reference structure, we estimate that NOE contacts had an average precision of 83% for the 12 modeling targets (**Figure S1b**). Importantly, only approximately half of the residues in the protein chain had at least one true positive (TP) NOE contact assigned (**Figure S1c**). RDC and □/ψ angle restraints were available for 43% and 49% of the residues on average. Surprisingly, the precision of the MetaPSICOV EC contact restraints (computed as fraction of residue-residue pairs with a Cβ-Cβ distance <8 Å among the top-L scoring |i-j| ≥ 6 contacts) was clearly better (93%) than for the simulated NOE restraints reflecting the great improvement in the computational contact prediction methods.

Our submitted models agreed with the NMR restraints nearly as good as the native structure. The number of NOE contacts made in the submitted Model 1 was comparable to the contacts made in the reference structure, and, importantly, a considerable number of TP-NOEs was made (**Table S1**). Excluding the two targets with the lowest GDT-TS score (N0989, N0981-D3) the average recall (i.e. number of TP-NOEs satisfied in the model vs. the native structure) was ca. 89%, and the average precision (i.e. number of TP-NOEs vs. number of all satisfied NOEs in the restraint set) was 86%. In addition, Rosetta models showed good RDC Q-factors^40^ (**Table S2**); the average RDC Q-factor was 43% (excluding targets N0989 and N0981-D3).

Model accuracy was significantly improved in regions with rich NOE contact information, and a clear inverse correlation between model-to-reference structure Cα-atom distance deviation and number and location of TP-NOE restraints was observed (see **Figure S2** and **S3**). However, we also found cases with accurate backbone structure predictions despite limited or missing NOE contact information (e.g. for targets N0968s1 and N1008) showing that Rosetta modeling can supplement sparse data. Importantly, modeling was not misguided by erroneous NOE data; no significant correlation between model accuracy and number and location of FP-NOEs could be found.

### Accuracy of Rosetta models and comparison to other methods

Our modeling strategy consisted of a two-stage approach (**Figure *1***): fold-level modeling with RosettaAbInitio and structure refinement with RosettaCM. RosettaAbInitio calculations rarely arrived at accurate models; the average GDT-TS score of the ten best scoring models over all targets was 30.5. Enhanced structural resampling via iterative model hybridization and fragment insertion with the RosettaCM protocol proved very effective in refining initial RosettaAbInitio models. Starting from these, the GDT-TS improved for all 12 targets, in some cases significantly (N0981-D5, ΔGDT-TS = 36.3). The average increase in GDT-TS over all targets was 16.7. **Figure S4** summarizes the improvement in GDT-TS and shows our submitted models for all targets.

Model accuracy in CASP13 was assessed on the level of the full-length protein as well as individual domains yielding 16 evaluation units for the investigated 12 monomeric targets. From our first submitted *de novo* models built with NOEs and RDCs, six out of 16 evaluation units had GDT-TS scores >60 and nine had GDT-TS scores >45 (see **Figure *2*a**), at which typically the native fold is correctly predicted (TM-score >0.5)^41^. Using exclusively RDCs as restraints for model folding and refinement was insufficient to yield accurate models. Only for 1/16 and 5/16 evaluation units the GDT-TS score was >60 and >45, respectively (**Figure *2*a**).

**Figure 2.**
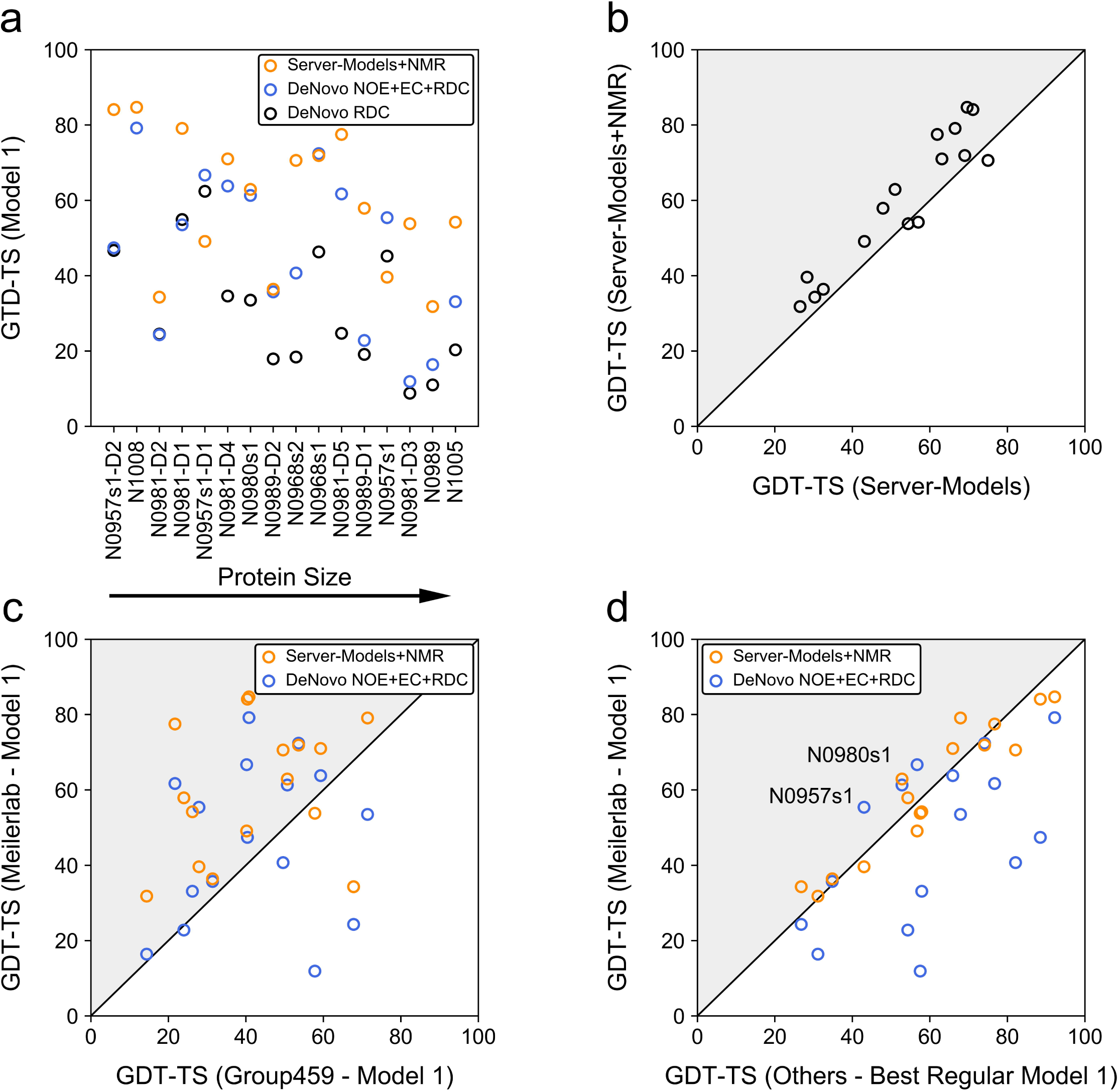
Overview of model GDT-TS of Meilerlab models obtained with different restraint sets and comparison to other structure predictions in the NMR-assisted and free modeling category. **(a)** GDT-TS of *de novo* predicted models created in CASP13 using only RDCs or NOEs, ECs and RDCs, respectively. In addition, the GDT-TS obtained when modeling started from server-models and employed an iterative refinement with NOEs, ECs, and RDCs, which was investigated in our post-CASP13 analysis, is shown. **(b)** Improvement of server-models through refinement with NMR data. The GDT-TS of the best model among all submissions of five different servers (Robetta^34^, I-TASSER^35,36^, QUARK^37^, RaptorX-Contact^38^, RaptorX-TBM^39^) (x-axis) is compared to the GDT-TS of our best-scoring model after hybridization and NMR-refinement of the respective server-models (y-axis). **(c)** Comparison of GDT-TS of the Meilerlab submitted Model 1 and NMR-refined server-models to Model 1 created by the ‘baseline’ group (Group 459) in the NMR-assisted category. Gray triangles indicate an improvement of the GDT-TS. **(d)** Comparison of GDT-TS of the Meilerlab submitted Model 1 and NMR-refined server-models to the best Model 1 in the regular unassisted modeling category.

Because the accuracy of our RosettaAbInitio starting models was generally low (only for three targets the GDT-TS was >45), we explored in our post-CASP13 analysis a meta-approach and tested whether the use of server-models as templates for structure refinement with RosettaCM would have improved our predictions. To this end, we chose the five submitted models of the Robetta^34^, I-TASSER^35,36^, QUARK^37^, RaptorX-Contact^38^ and RaptorX-TBM^39^ server, and hybridized and refined those models through 20 rounds of RosettaCM. The model with the lowest score after the last refinement step was deemed the final model. With this procedure, the number of predictions with GDT-TS >60 and >45 increased to eight and twelve out of 16 evaluation units (**Figure *2*a**). Furthermore, the NMR restraints helped to consistently improve model accuracy, and for 13/16 evaluation units the GDT-TS score after NMR-restrained Rosetta refinement was higher than the GDT-TS of the best model among the original 25 server-models (**Figure *2*b** and **Figure S5**).

Comparing our approach with other methods, we find that Rosetta calculations produced more predictions with higher GDT-TS score than the ‘baseline’ method in the NMR-assisted category (a hybrid method of ASDP^42^ and CYANA^43,44^) operating on the same restraint set (NOEs, RDCs and ECs). In 11/16 and 14/16 cases, our NMR-restrained *de novo* models and NMR-refined server-models, respectively, had higher GDT-TS scores (compare with **Figure *2*c**). However, it is a surprising result that our first submitted models are often not better than the best models from the non-assisted free modeling category (**Figure *2*d**). This may reflect the big advancement in contact prediction methods which have pushed protein *de novo* structure prediction forward. In CASP13, the three contact prediction methods with the overall best performance achieved for this particular set of protein targets presented here (excluding N0981-D1, N0981-D4 and N0981-D5 for which no predictions had been made), an average precision of 55%, 73% and 89%, respectively, evaluated on the top L, L/2 and L/5 long-(|i-j| ≥ 24) and medium-range (12 ≤ |i-j| ≤ 23) contacts. The difference in model GDT-TS becomes more balanced when the free modeling predictions are compared to our NMR-refined server-models which shows that with adjustments to the protocol Rosetta can achieve equal performance.

Examples of models for which high accuracy was achieved are shown in **Figure 3a-c**. The Cα-RMSD from the native structure of the first submitted model for these three targets was below 3.5 Å (N1008: 2.06 Å, N0968s1: 2.88 Å, N0981-D5: 3.48 Å) enabling accurate sidechain placement. Furthermore, we submitted the best prediction for target N0957s1 (GDT-TS = 56.0) among all predictors in the data-assisted and free-modeling categories.

**Figure 3.**
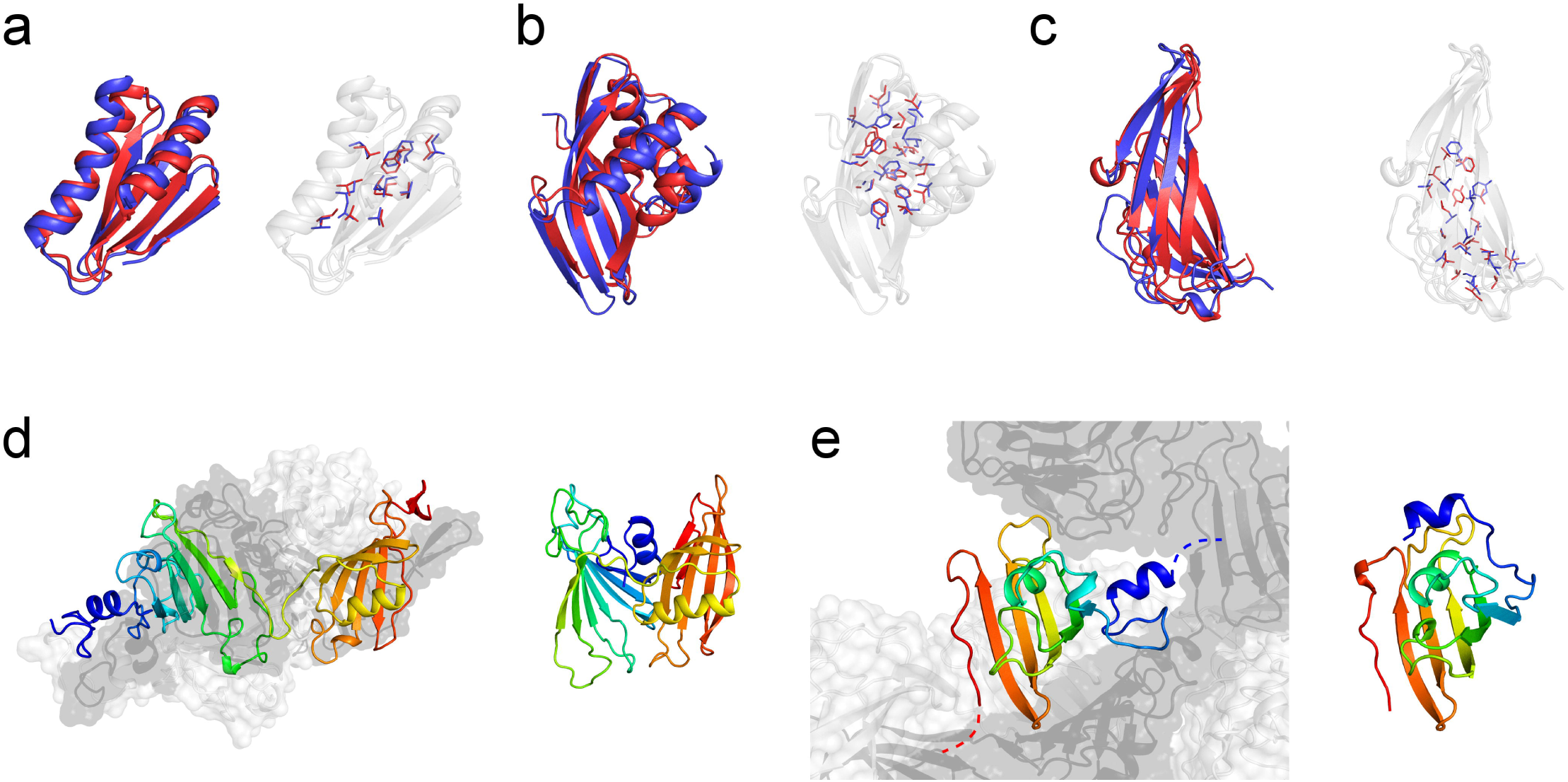
Examples of successful and difficult modeling cases. High-resolution structure predictions could be made for three targets, **(a)** N1008 (Cα-RMSD = 2.06 Å), **(b)** N0968s1 (Cα-RMSD = 2.88 Å), and **(c)** N0981-D5 (Cα-RMSD = 3.48 Å), allowing accurate sidechain placement. The submitted Model 1 (red) is displayed as cartoon representation and compared with the experimental reference structure (blue). Sidechains in the protein core are depicted with sticks. The fraction of correct χ_1_ and χ_2_ rotamers in buried protein regions was **(a)** 74% / 37%, **(b)** 77% / 41%, and **(c)** 66% / 29%, respectively. Difficult modeling cases were proteins with quaternary structure which have interfaces to other protein copies or domains. **(d)** Target N0989 is a homo-trimeric complex. The monomer was modeled with a too compact, non-extended conformation. **(e)** The model of target N0981-D4 shows an incorrect orientation of its N-terminal helix (blue) which is linked to another domain in the native assembly. No NOE contact information was available for this part of the structure.

### What went wrong: Importance of quaternary modeling and model bias from missing NMR data

Examples of unsuccessful structure predictions are given in **Figure 3d+e**. These include proteins with quaternary structure in which the target represents one subunit. Because of time limitations we did not pursue quaternary modeling for homo-oligomeric targets which is a notoriously difficult task when starting from *de novo* models and when interdomain contact information is insufficient. However, modeling of the oligomeric state becomes important when the protein is stabilized by interactions with neighboring subunits. This is the case for target N0989 where the N- and C-terminal domains make very little contacts with each other but are packed towards chain B and C (**Figure 3d**). Consequently, the Rosetta model of the monomer was predicted with a too compact, non-elongated conformation. The second example, N0981-D4, features an extended N-terminal helix that connects to the next domain and is stabilized by quaternary interactions. In our Rosetta model of the single domain (**Figure 3e**), the N-terminal helix is folded onto the central β-sheet to maximize residue burial and because this helix was not restrained by NOE contact information.

For hetero-dimeric target N0980 modeling of the oligomeric state was carried out with RosettaDock starting from our *de novo* predictions of the two separate domains. This strategy turned out to be suboptimal because the small domain (N0980s2, chain B) was incorrectly modeled as compact globular protein, but adopts an elongated conformation wrapping around the larger domain (N0980s1, chain A) in the native structure. The ligand RMSD (computed on chain B after superimposition of chain A) of our submitted model was 22.7 Å. A more suitable modeling strategy may involve simultaneous folding and docking of the smaller protein domain onto the larger domain which we explored in our post-CASP13 analysis and which improved the ligand RMSD up to 12.8 Å (**Figure S6**).

The remaining targets with low model accuracy (N0981-D1, N0981-D2, N0981-D3) had poor quality fragments, especially N0981-D3 which had no 9mer fragment <1 Å to the native structure for 75% of its residues (compare with **Figure S7**). This target was difficult in several aspects. It had little regular secondary structure (≤46%) and an unusual β-sandwich topology with adjacent strands switching back and forth between two β-sheets.

Model accuracy of some targets suffered from missing NOE contact information leading to wrong domain orientations or loop conformations (compare with **Figure S2** and **S3**). Examples of those regions in our submitted models include the long C-terminal loop in N0980s1, the N-terminal helix in N0981-D4 and the C-terminal β-sheet in N0968s2 which was flipped upside down. However, we also find instances in which high accuracy was achieved in regions having no or very little NOE contacts. For example, target N1008 used the fewest NOE contacts (0.65 per residue) with the lowest precision (~60%) but was predicted with 2.06 Å Cα-RMSD to the native structure.

## Discussion

NMR data as limited as around two non-local TP-NOEs per residue and less than one RDC and dihedral restraint per residue were sufficient to generate protein models with the correct fold. In some cases, even high-resolution models were achieved with Cα-RMSD better than 3.5 Å from the native structure. Model accuracy was clearly superior in regions with more NOE contacts indicating that Rosetta can translate the contact information in biasing the model to the native structure.

Enhanced structural sampling which was accomplished in this study by iterative model hybridization and fragment insertion with RosettaCM was crucial to leverage the NMR restraints and improve model quality. Refinement made use of ambiguous contact information leading to significant improvements (~17 GDT-TS units on average) over RosettaAbInitio models generated with only non-ambiguous contacts. Use of template information in the form of server-models as input to NMR-guided RosettaCM refinement led to an additional increase in the average model GDT-TS score from 46.6 to 59.9 as observed in our post-CASP13 analysis. This amounts to half of the targets being modeled with higher accuracy than the best free modeling predictions. Encouraged by this result, we believe that a combination of NMR data with orthogonal structural information derived from e.g. template structures and predicted contacts (as outlined in **Figure 1c**) will be an important new driving force to make constructive progress in NMR structure prediction. This includes new areas of applications such as NMR-guided detection of templates^11^ or the use of ECs in assisting NMR data assignment/interpretation and NMR structure calculation^45^. In addition, computational models created by such meta-approaches will have the advantage that they can be experimentally validated e.g. by comparison against a set of NOE contacts hold out for cross-validation.

Additional bioinformatical or experimental contact restraints should be selected ideally from regions with incomplete or missing NMR assignment which would help restraining those parts of the model and could resolve wrong domain orientations as described above. Next CASP experiments could also introduce other types of NMR data. For example, ^1^H^N^, ^13^Cα, ^13^Cβ, ^13^C’ and ^15^N^H^ chemical shifts can be used directly in the fragment picking process which avoids errors in the backtranslation to torsion angle restraints and could improve fragment quality. Paramagnetic NMR data, e.g. pseudocontact shifts (PCSs), may be used as additional source of long-range structural restraints.

Subsequent CASP experiments could also explore the possibility to model multiple conformational states and conformational transitions in proteins – a challenge which has not been examined in the current CASP but to which NMR provides very sensitive tools of detection. For example, information on protein dynamics and a detailed description of the structure of alternative conformational states can be inferred from relaxation dispersion and chemical exchange saturation transfer data. In this scenario, NOE contacts may be used with greater caution because part of the contacts may be incompatible with each other.

## Supplemental Information

Supporting information includes seven additional figures and two additional tables which may be found in the online version of this article.

## Acknowledgements

The authors thank the members of the RosettaCommons for their helpful discussion and support, especially Dr. Sergey Ovchinnikov and Dr. Hahnbeom Park for their original development of the iterative RosettaCM refinement protocol. The authors would also like thank the CASP13 organizers, especially Dr. Montelione and the members of his group for preparing NMR data and assessing predictions, and the structural biologists who generously provided structure targets. This work was conducted using the computing cluster of the Advanced Center for Research and Education (ACCRE) at Vanderbilt University, Nashville, TN.

