## Supplemental Information for "Protein structure prediction using sparse NOE and RDC restraints with Rosetta in CASP13"

### Supporting Figures

#### Figure S1: Richness and quality of the used NMR datasets

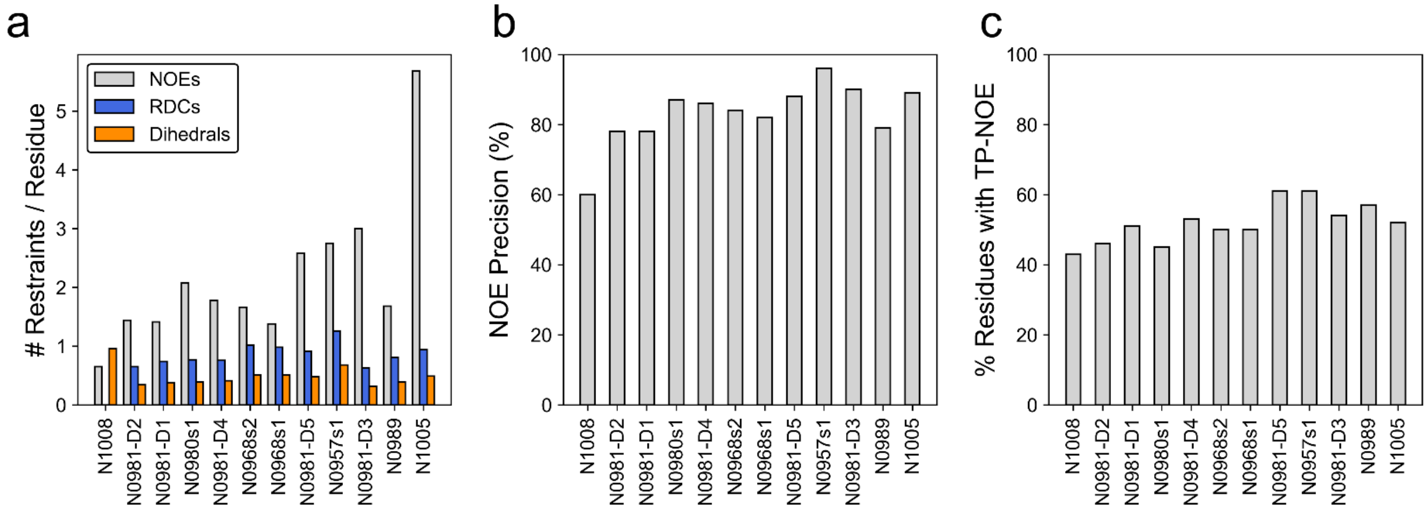

**Figure S1: Richness and quality of the used NMR datasets. (a)** Number of per-residue NOE, RDC and ϕ/ψ dihedral angle restraints which were employed in our NMR-assisted structure predictions. The average number of NOE, RDC and dihedral restraints was 2.2, 0.8 and 0.5 per residue, respectively. **(b)** NOE precision; defined as ratio of the number of true positive (TP)-NOEs to the total number of NOEs. The average NOE precision was 83%. **(c)** Fraction of protein residues which had at least one TP-NOE. The average over all prediction targets was 52%.

Figure S2: Local model accuracy of *de novo* predicted NMR-assisted targets

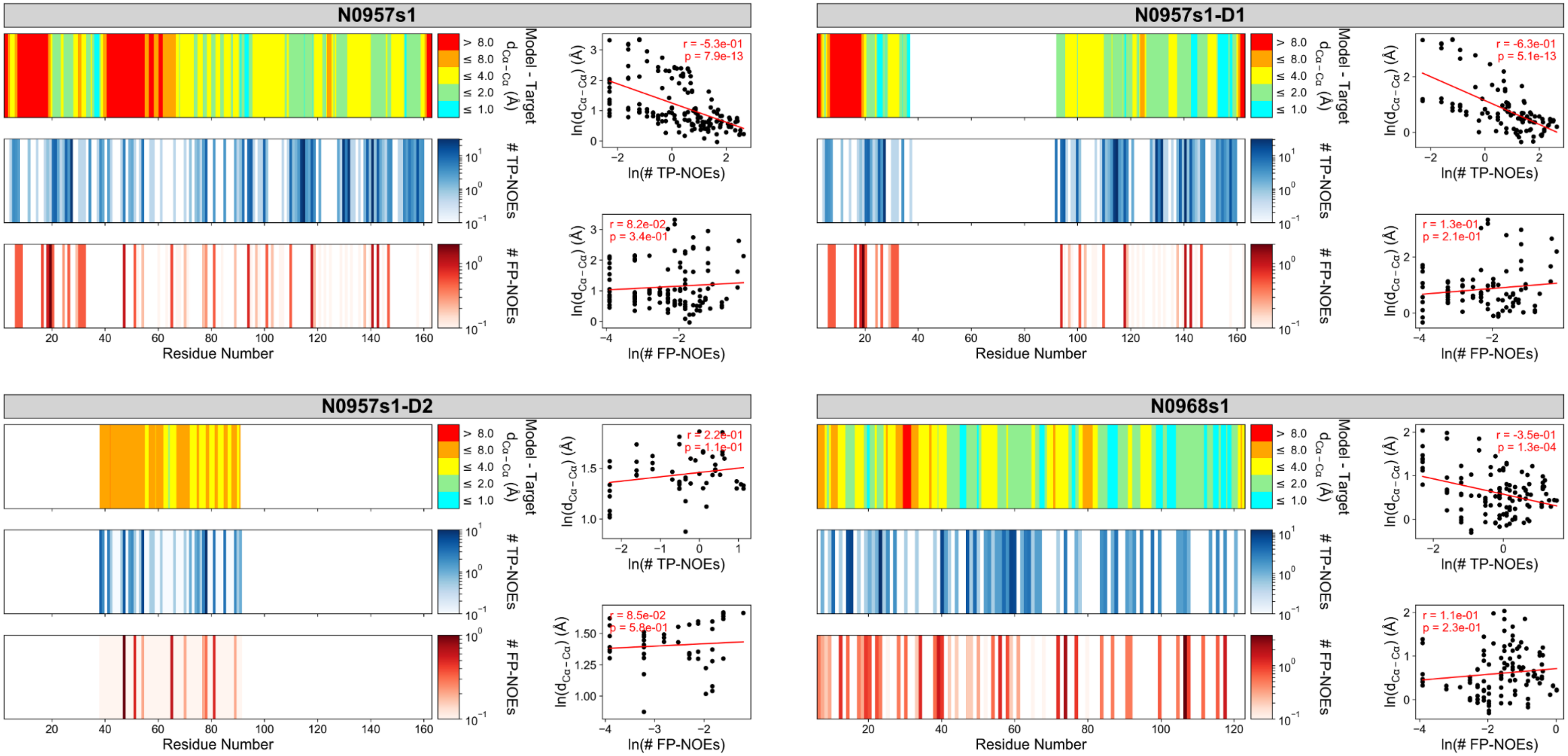

**Figure S2: Comparison of local model accuracy of *de novo* predicted NMR-assisted targets to number and distribution of NOE restraints.** For each of the 12 prediction targets or 16 evaluation units, respectively, the local Cα-Cα deviation between model and reference structure is compared to the number and location of true positive (TP) and false positive (FP) NOEs. For targets N0957s1 and N0989, data are also shown for separate domains. For each target, the panels display the following results: On the left side, the Cα-Cα distance between the submitted model 1 and the reference structure after superimposition with the program MAMMOTH[1] (top) and the number of TP (middle) and FP (bottom) NOEs versus the protein residue number are shown. On the right side, the natural logarithm of the Cα-Cα distance deviation (calculated as running average over five neighboring residues) is plotted versus the logarithm of the number of TP (top) and FP (bottom) NOEs. Red lines represent the least-squares fit to a linear regression model.

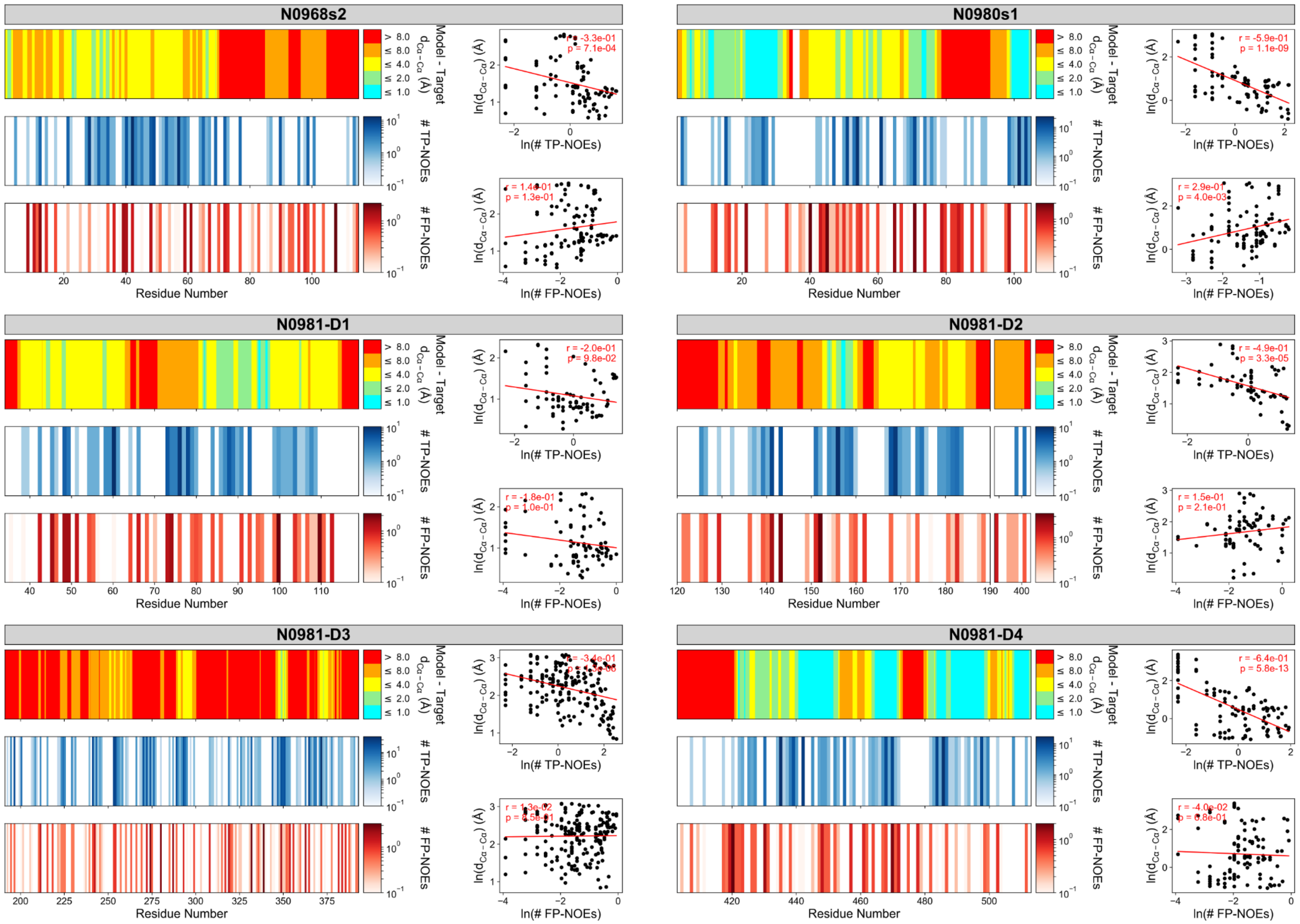

**Figure S2: Comparison of local model accuracy of *de novo* predicted NMR-assisted targets to number and distribution of NOE restraints. (continued)**

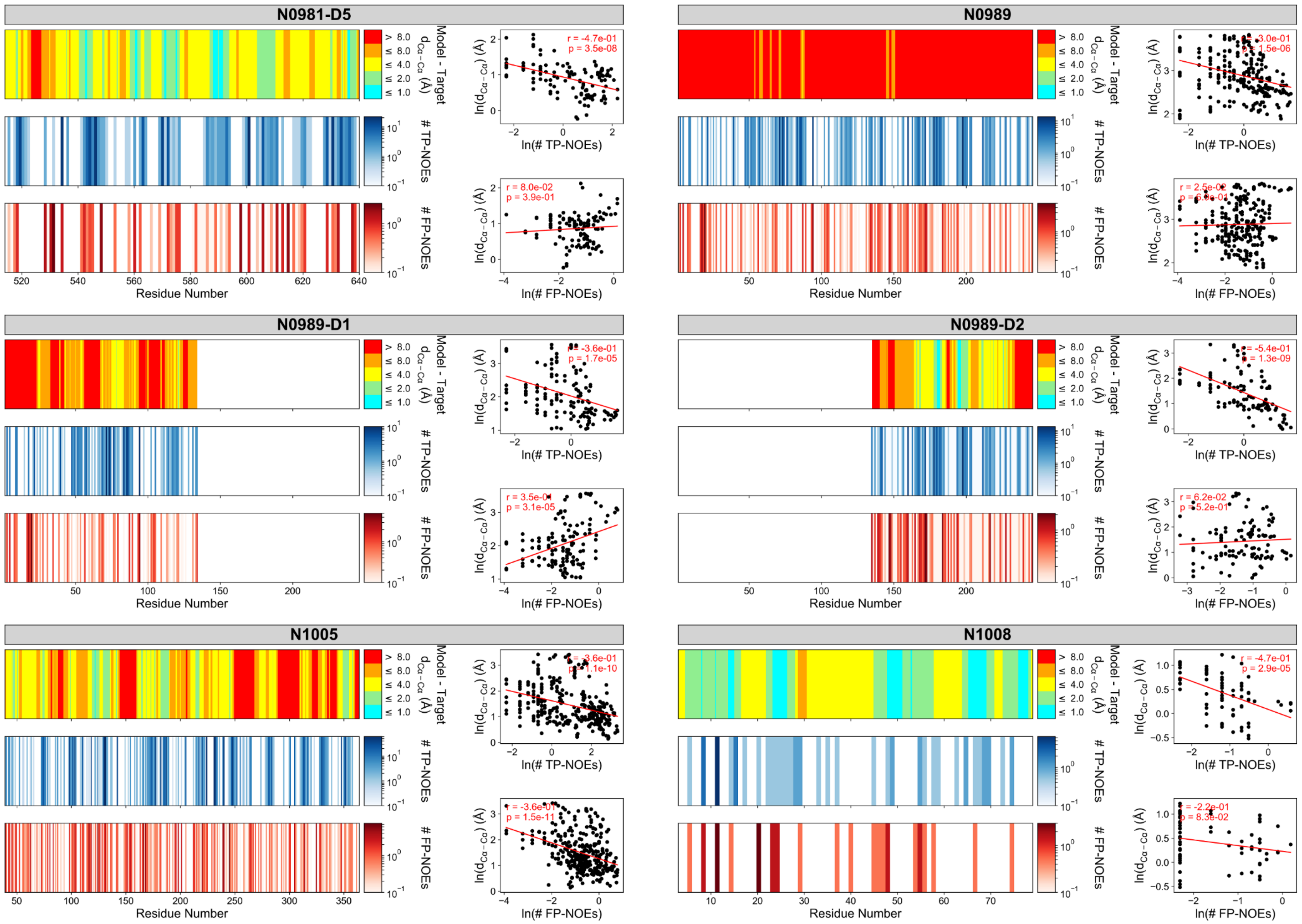

**Figure S2: Comparison of local model accuracy of *de novo* predicted NMR-assisted targets to number and distribution of NOE restraints. (continued)**

#### Figure S3: Local model accuracy of NMR-refined server-models from our post-CASP13 analysis

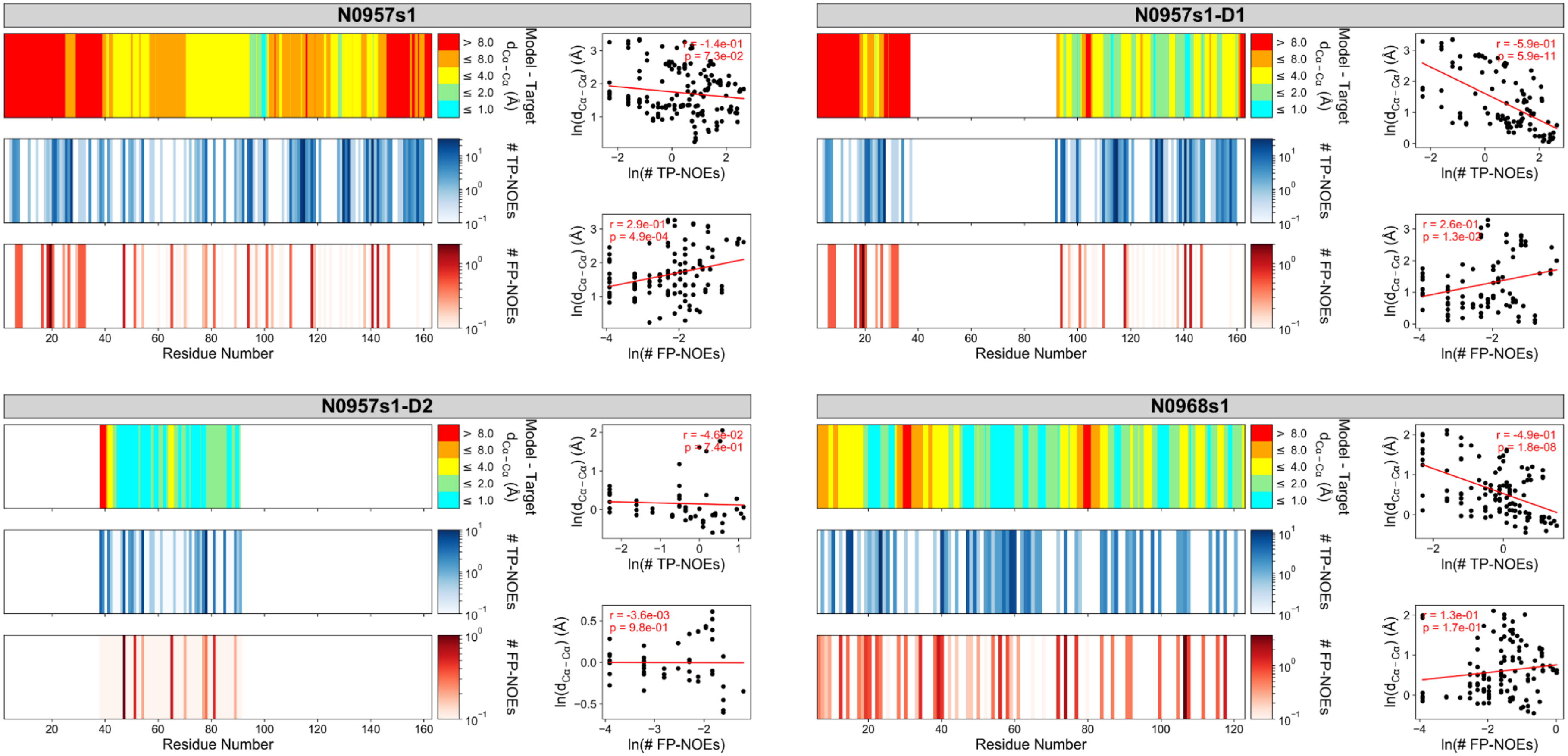

**Figure S3: Comparison of local model accuracy of NMR-refined server-models from our post-CASP13 analysis to distribution and number of NOE restraints.** Models were created by combining and refining the five submitted models from the I-TASSER[2,3], QUARK[4], Robetta[5], RaptorX-Contact[6] and RaptorX-TBM[7] server, respectively, with RosettaCM and NMR data. For each of the 12 prediction targets or 16 evaluation units, respectively, the local Cα-Cα deviation between the best scoring NMR-refined server-model and the reference structure is compared to the number and location of true positive (TP) and false positive (FP) NOEs. For targets N0957s1 and N0989, data are also shown for separate domains. The arrangement of panels is the same as in **Figure S2**: On the left side, the Cα-Cα distance between model and reference structure after superimposition with the program MAMMOTH[1] (top) and the number of TP (middle) and FP (bottom) NOEs versus the protein residue number are shown. On the right side, the natural logarithm of the Cα-Cα distance deviation (calculated as running average over five neighboring residues) is plotted versus the logarithm of the number of TP (top) and FP (bottom) NOEs. Red lines represent the least-squares fit to a linear regression model.

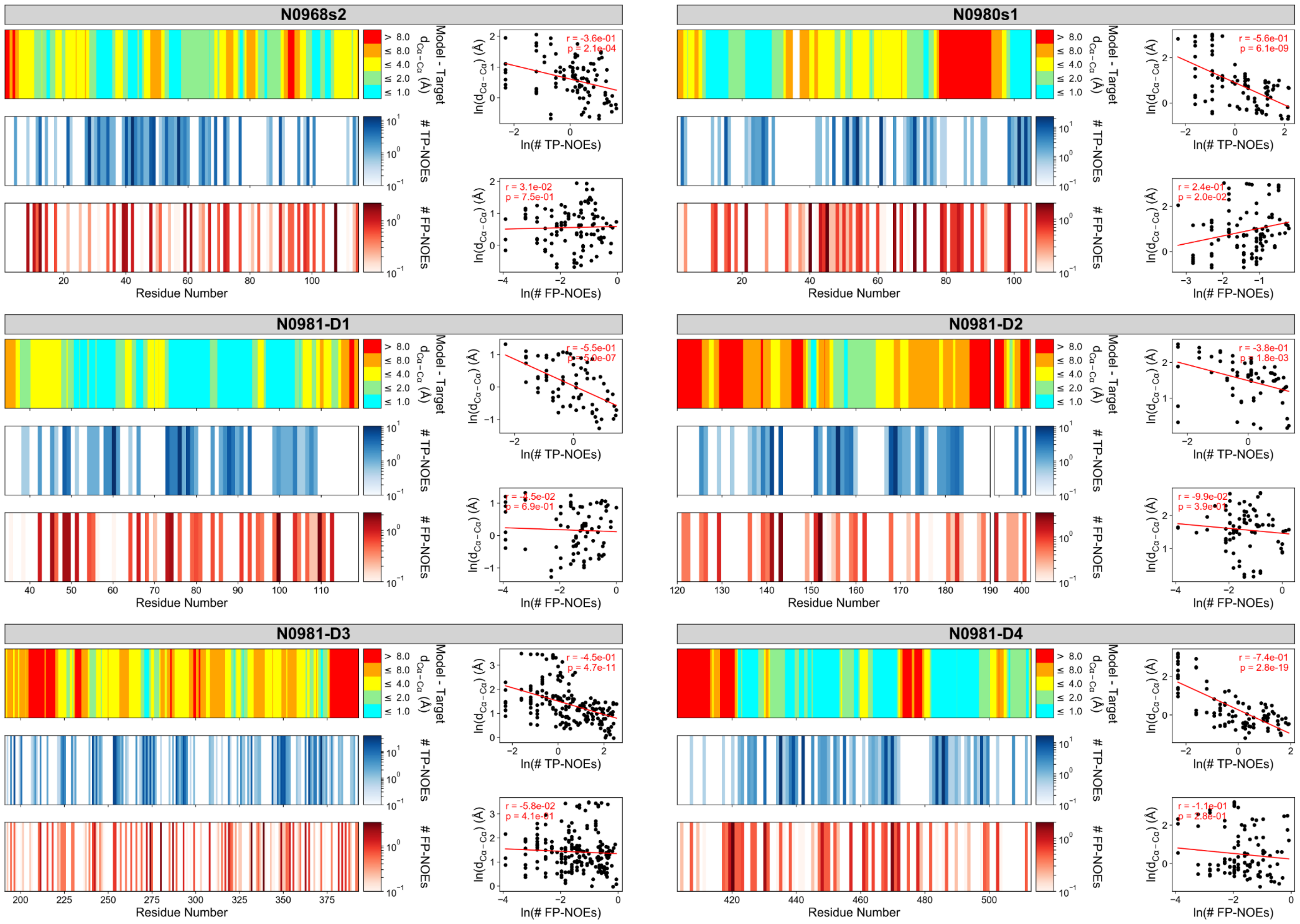

**Figure S3: Comparison of local model accuracy of NMR-refined server-models from our post-CASP13 analysis to distribution and number of NOE restraints (continued).**

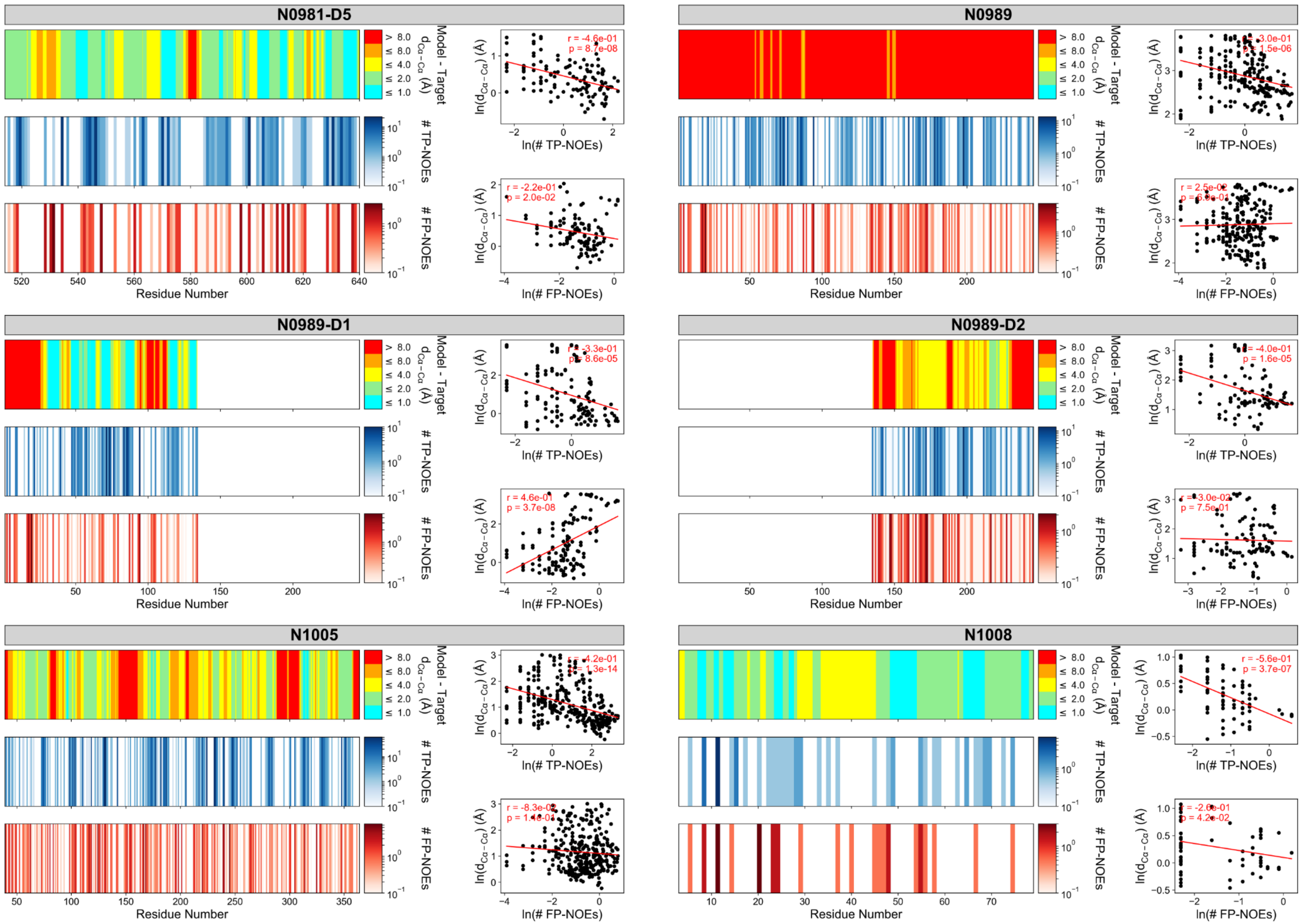

**Figure S3: Comparison of local model accuracy of NMR-refined server-models from our post-CASP13 analysis to distribution and number of NOE restraints (continued).**

Figure S4: Results of *de novo* structure prediction of NMR-assisted targets

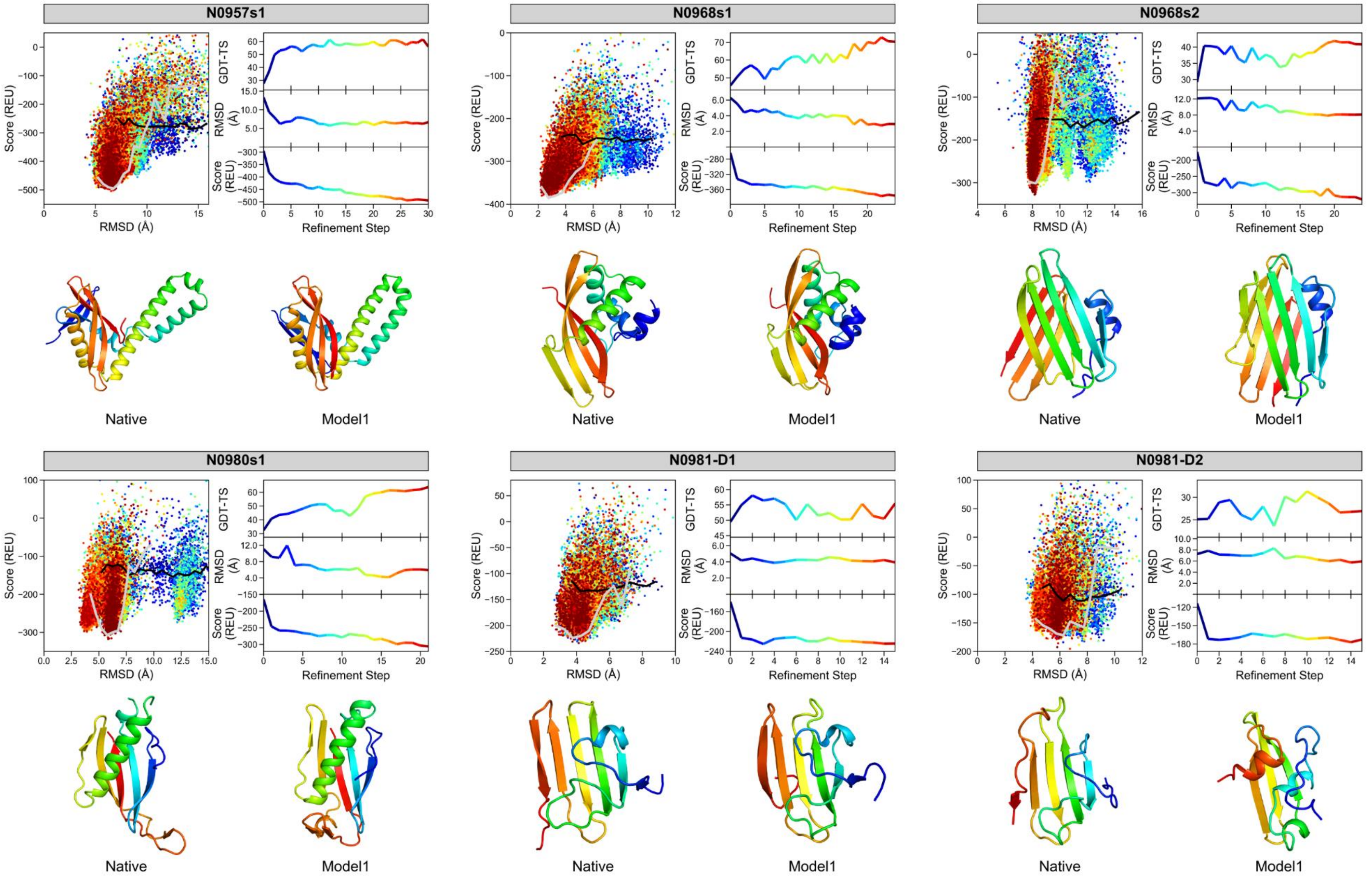

**Figure S4: Results of *de novo* structure prediction of NMR-assisted (NOE+RDC) modeling targets in CASP13.**

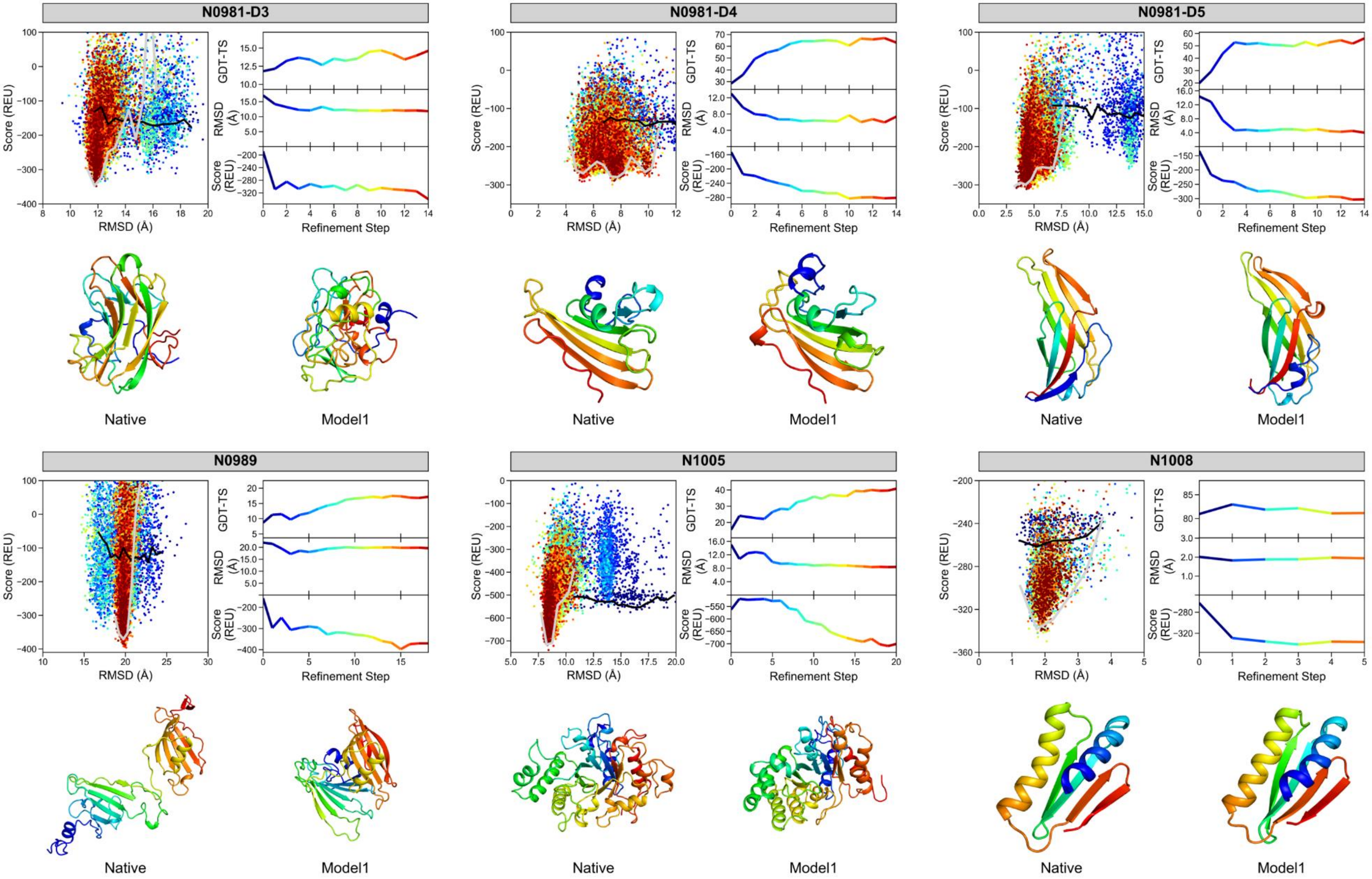

**Figure S4: Results of *de novo* structure prediction of NMR-assisted (NOE+RDC) modeling targets in CASP13 (continued).** Each of the 12 prediction targets was first folded with RosettaAbInitio and then iteratively refined with RosettaCM. The panels display the following results: The upper left panel plots the combined Rosetta energy and NMR NOE+RDC restraint score versus the model's Cα-RMSD relative to the reference structure. The color coding (blue to red) corresponds to the number of refinement steps. The black and gray line represent the lowest-energy rim (i.e. the median of the five lowest-scoring models by RMSD bin) of the score-vs-RMSD plot of the model pool before the first and after the final refinement step. Note that the pool at step 0 is comprised of models after the initial iterative hybridize selection step. The upper right panel displays the average GDT-TS (top row), Cα-RMSD (middle row) and score (bottom row) of the ten lowest scoring models after each refinement step. The values of successive refinement steps are colored from blue to red. The lower panel compares the experimental reference structure (left) depicted as ribbon diagram and colored in rainbow with the submitted model 1 (right) which was the best-scoring model after the last refinement step.

### Figure S5: Results of NMR-assisted refinement of server-models in our post-CASP13 analysis

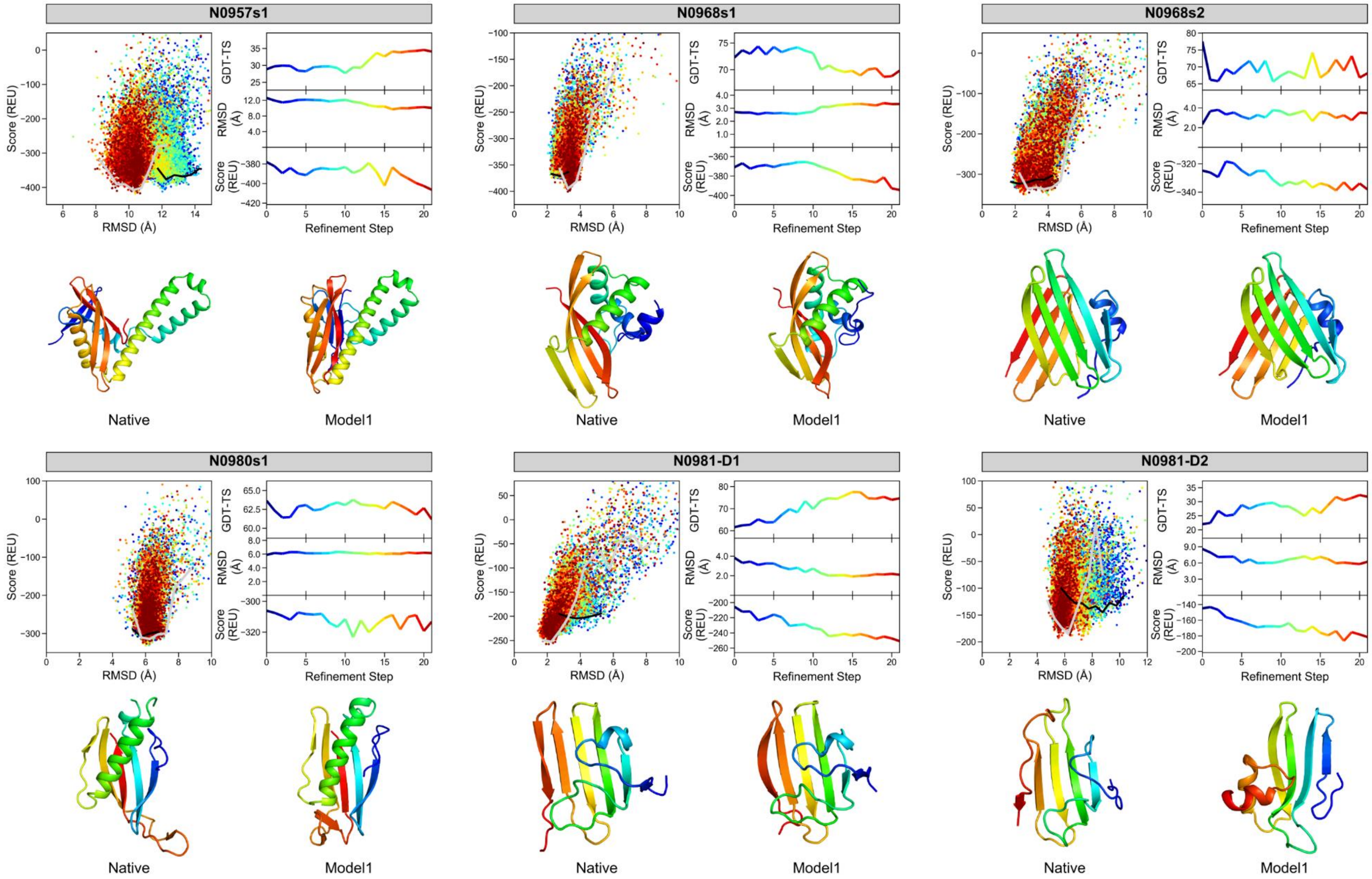

**Figure S5: Results of NMR-assisted structure refinement of server-models in our post-CASP13 analysis.**

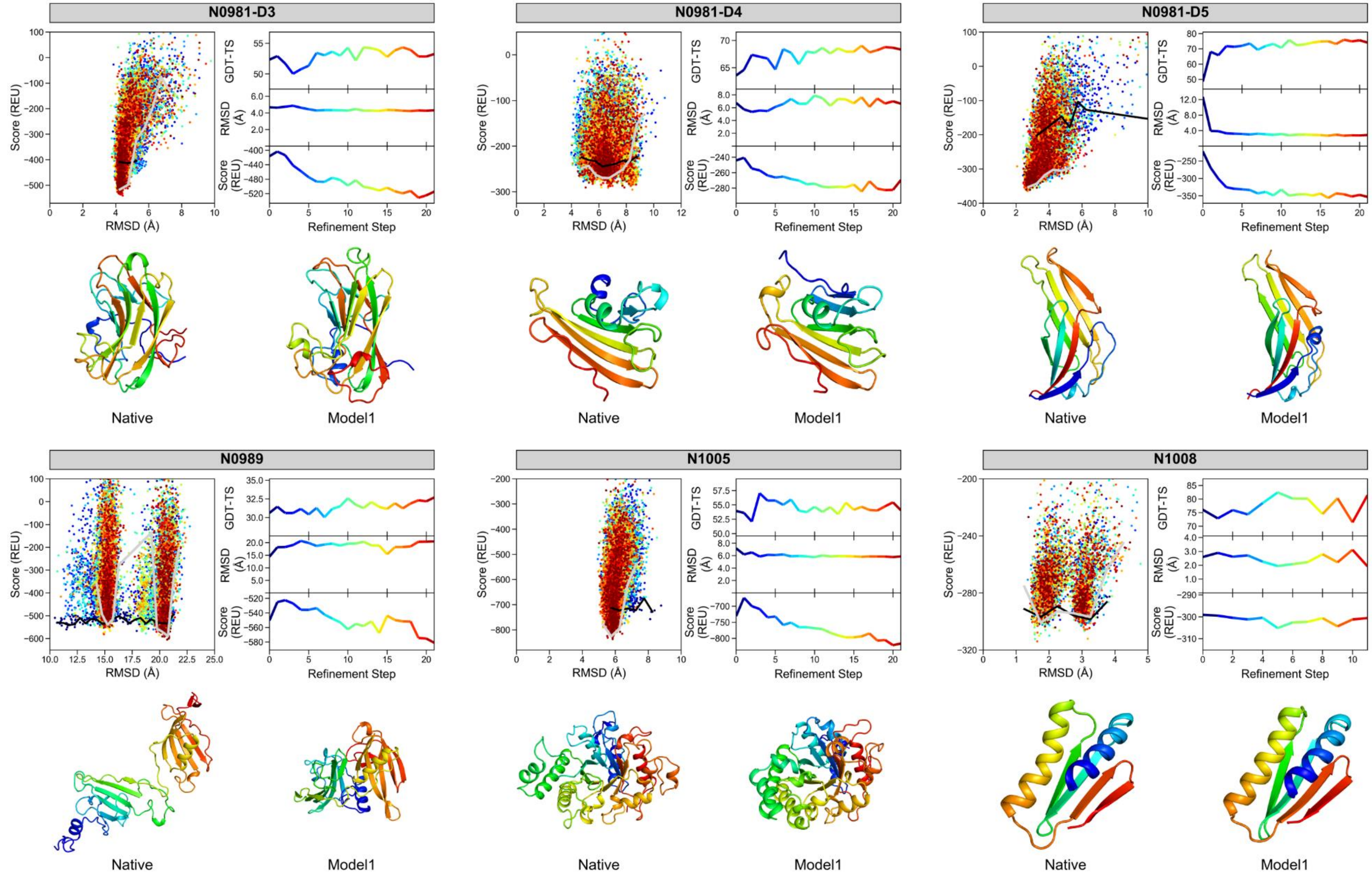

**Figure S5: Results of NMR-assisted structure refinement of server-models in our post-CASP13 analysis (continued).** For each of the 12 prediction targets, the top five models submitted by the I-TASSER[2,3], QUARK[4], Robetta[5], RaptorX-Contact[6] and RaptorX-TBM[7] server were combined and iteratively refined with RosettaCM and NMR (NOE+RDC) data. The arrangement of panels and the type of displayed structure evaluation metrics are the same as in **Figure S4**: The upper left panel plots the combined Rosetta energy and NMR NOE+RDC restraint score versus the model's Cα-RMSD relative to the reference structure. The upper right panel displays the average GDT-TS (top row), Cα-RMSD (middle row) and score (bottom row) of the then lowest scoring models after each refinement step. The values of successive refinement steps are colored from blue to red. The lower panel compares the experimental reference structure (left) depicted as ribbon diagram and colored in rainbow with the lowest scoring model after the last refinement step (right).

Figure S6: Structure prediction of oligomeric target H0980

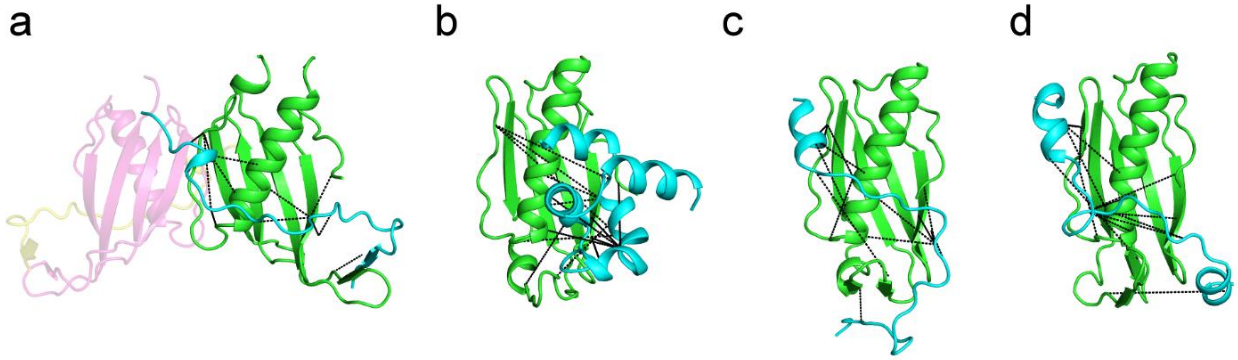

**Figure S6: Structure prediction of oligomeric target H0980. (a)** The asymmetric unit of the X-ray structure contains two copies of the receptor (colored green and pale magenta) and peptide (colored cyan and pale yellow) chain, displayed in cartoon representation. **(b)** Our submitted model 1 was created by (i) folding the receptor and peptide chain separately and (ii) docking both models together with RosettaDock[8] guided by intermolecular NOE distance restraints (depicted as black dotted lines) and RDCs. The peptide ligand Cα-RMSD after superimposition of the receptor was 22.7 Å. An alternative modeling strategy that we explored in our post-CASP13 analysis involved simultaneous folding and docking of the peptide onto the receptor model using Rosetta FlexPepDock[9]. The lowest-scoring FlexPepDock model **(c)** and the lowest-RMSD model **(d)** had a ligand Cα-RMSD of 16.0 Å and 12.8 Å, respectively.

#### Figure S7: Local model accuracy and fragment quality of *de novo* predicted NMR-assisted targets

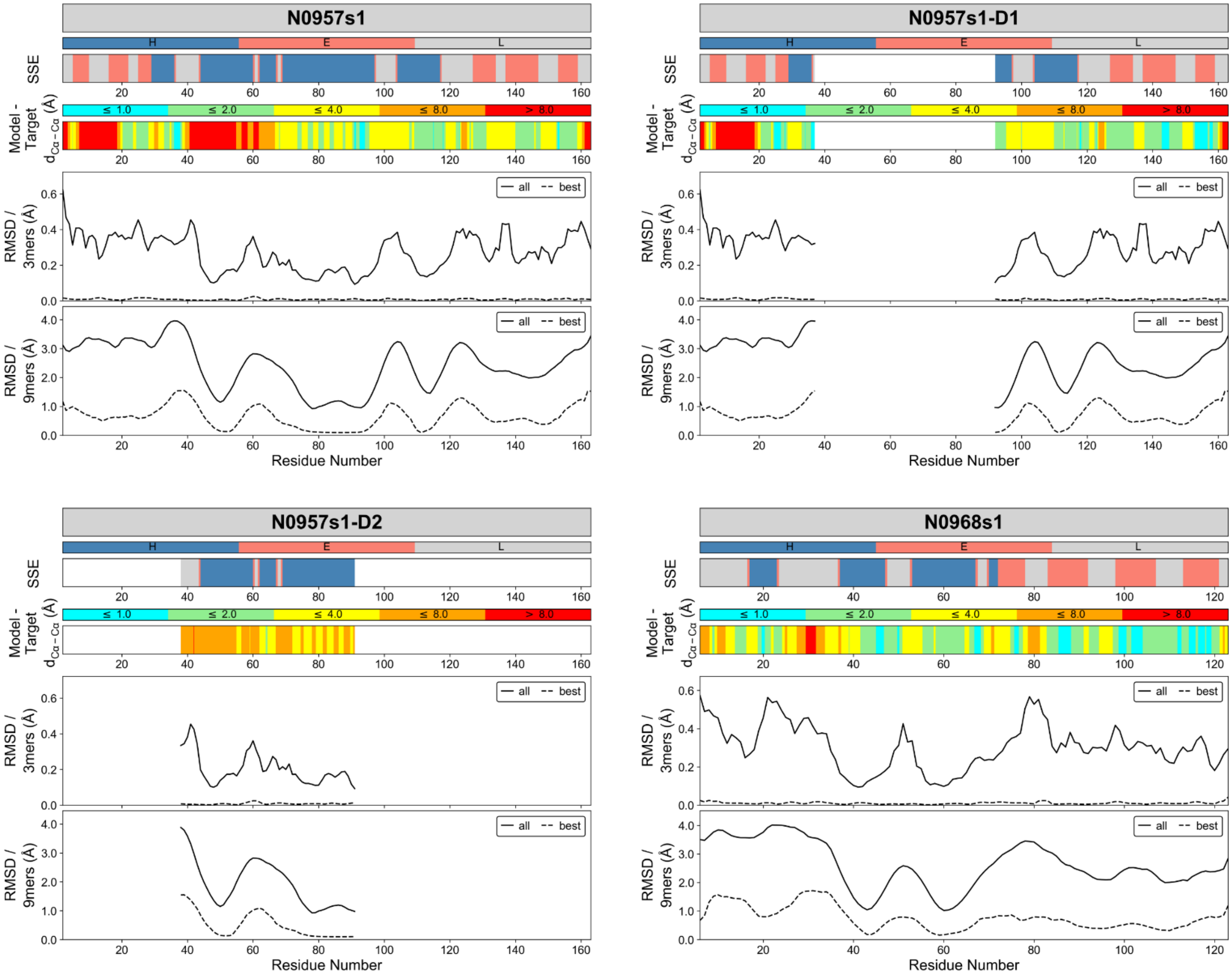

**Figure S7: Comparison of local model accuracy of *de novo* predicted NMR-assisted (NOEs+RDCs) targets to Rosetta fragment quality.** For each of the 12 prediction targets or 16 evaluation units, respectively, the local Cα-Cα deviation between model and reference structure is compared to the Cα-RMSD of the employed 3- and 9-residue fragments and the target secondary structure. For each target the panels display the following information: DSSP[10]-assigned secondary structure (top panel; blue: helix, salmon: sheet, gray: loop), model-target Cα-Cα distance deviation (second panel from top; cyan: ≤1Å, green: ≤2Å, yellow: ≤4Å, orange: ≤8Å, red: >8Å), Cα-RMSD of 3-residue fragments along protein sequence (second panel from bottom; solid line represents average over all 200 fragments and broken line represents lowest RMSD fragment) and Cα-RMSD of 9-residue fragments along protein sequence (bottom panel; solid line: average RMSD all fragments, broken line: lowest RMSD fragment).

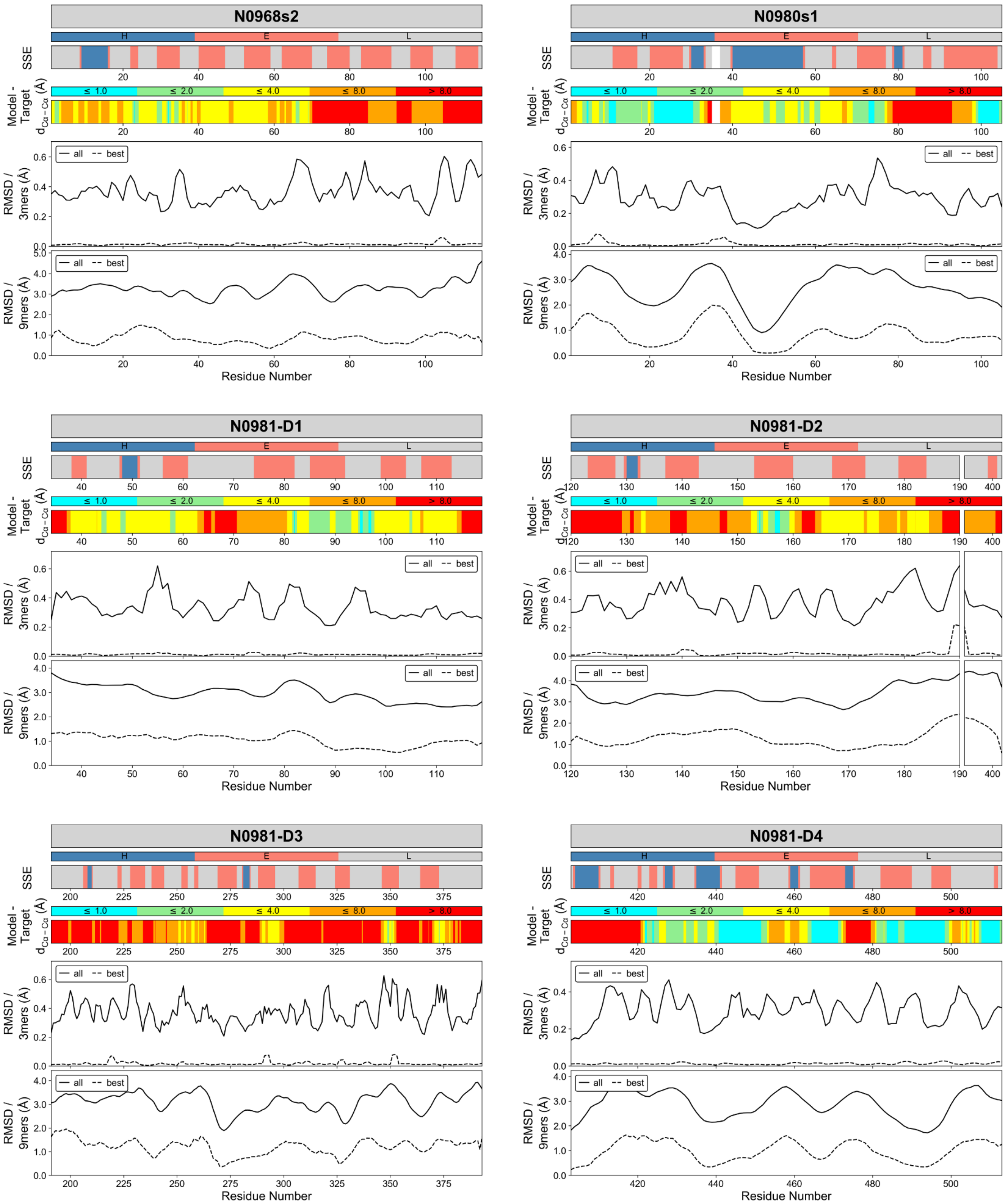

**Figure S7: Comparison of local model accuracy of *de novo* predicted NMR-assisted (NOEs+RDCs) targets with Rosetta fragment quality (continued).**

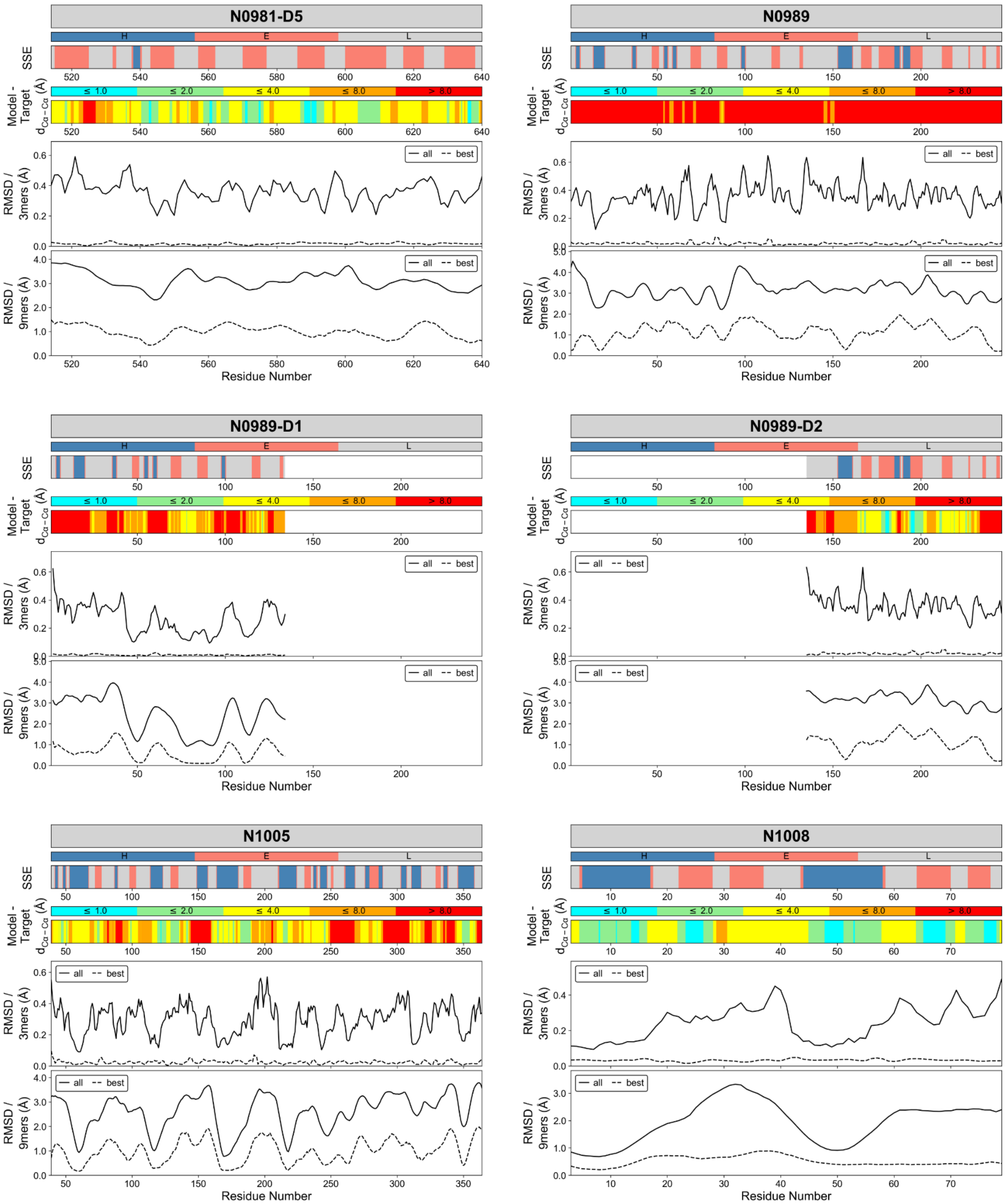

**Figure S7: Comparison of local model accuracy of *de novo* predicted NMR-assisted (NOEs+RDCs) targets with Rosetta fragment quality (continued).**

### Supporting Tables

#### Table S1: Satisfied NOE contacts in submitted *de novo* models and NMR-refined server-models

| Target | Residues | Reference Structure | | DeNovo+NOE+RDC Submitted Model 1 | | | | NMR-refined Server-Model | | | |
|---|---|---|---|---|---|---|---|---|---|---|---|
| | | All# | TP* | All# | TP* | Recall (%)† | Precision (%)‡ | All# | TP* | Recall (%)† | Precision (%)‡ |
| N1008 | 77 | 30 | 30 | 33 | 29 | 96.7 | 87.9 | 33 | 29 | 96.7 | 87.9 |
| N0981-D2 | 80 | 90 | 90 | 86 | 65 | 72.2 | 75.6 | 87 | 66 | 73.3 | 75.9 |
| N0981-D1 | 86 | 95 | 95 | 91 | 83 | 87.4 | 91.2 | 99 | 93 | 97.9 | 93.9 |
| N0980s1 | 111 | 200 | 200 | 231 | 193 | 96.5 | 83.5 | 233 | 194 | 97.0 | 83.3 |
| N0981-D4 | 111 | 171 | 171 | 184 | 166 | 97.1 | 90.2 | 185 | 166 | 97.1 | 89.7 |
| N0968s2 | 116 | 161 | 161 | 180 | 136 | 84.5 | 75.6 | 168 | 156 | 96.9 | 92.9 |
| N0968s1 | 126 | 143 | 143 | 142 | 139 | 97.2 | 97.9 | 144 | 142 | 99.3 | 98.6 |
| N0981-D5 | 127 | 288 | 288 | 278 | 250 | 86.8 | 89.9 | 292 | 271 | 94.1 | 92.8 |
| N0957s1 | 163 | 428 | 428 | 450 | 387 | 90.4 | 86.0 | 396 | 344 | 80.4 | 86.9 |
| N0981-D3 | 203 | 546 | 546 | 406 | 255 | 46.7 | 62.8 | 452 | 361 | 66.1 | 79.9 |
| N0989 | 246 | 327 | 327 | 290 | 190 | 58.1 | 65.5 | 381 | 273 | 83.5 | 71.7 |
| N1005 | 326 | 1640 | 1640 | 1555 | 1285 | 78.4 | 82.6 | 1668 | 1439 | 87.7 | 86.3 |
| **Mean** | | | | | | **82.7** | **82.4** | | | **89.2** | **86.6** |
| **Mean (-N0981-D3, -N0989)** | | | | | | **88.7** | **86.0** | | | **92.0** | **88.8** |

#Number of all satisfied NOE contacts in restraint set. For the native reference structure all satisfied NOE contacts are considered TP-NOEs.
*Number of satisfied TP-NOE contacts in restraint set.
†Recall is ratio of TP-NOE contacts in Rosetta model vs. TP-NOE contacts in reference structure.
‡Precision is ratio of satisfied TP-NOE contacts vs. all satisfied NOE contacts in Rosetta model.

#### Table S2: RDC Q-factor (%) of submitted *de novo* models and NMR-refined server-models

| Target | Residues | DeNovo+RDC submitted Model 1 | DeNovo+NOE+RDC submitted Model 1 | NMR-refined Server-Model |
|---|---|---|---|---|
| N1008* | 77 | - | - | - |
| N0981-D2 | 80 | 31.2 | 38.8 | 44.6 |
| N0981-D1 | 86 | 37.0 | 30.1 | 28.9 |
| N0980s1 | 111 | 45.3 | 36.7 | 32.8 |
| N0981-D4 | 111 | 57.5 | 51.3 | 62.2 |
| N0968s2 | 116 | 38.9 | 33.2 | 25.4 |
| N0968s1 | 126 | 40.7 | 37.5 | 36.3 |
| N0981-D5 | 127 | 55.0 | 51.2 | 30.7 |
| N0957s1 | 163 | 44.4 | 42.4 | 53.4 |
| N0981-D3 | 203 | 67.9 | 84.8 | 60.8 |
| N0989 | 246 | 52.7 | 49.4 | 38.3 |
| N1005 | 326 | 72.8 | 66.6 | 56.8 |
| **Mean** | | **49.4** | **47.4** | **42.7** |
| **Mean (-N0981-D3, -N0989)** | | **47.0** | **43.1** | **41.2** |

*No RDC data were available for target N1008
